# Selonsertib-Eluting Electrode Coating Attenuates Cochlear Injury Pathways

**DOI:** 10.64898/2026.04.02.716022

**Authors:** Jacqueline M Ogier, Lilith M Caballero-Aguilar, Michael G Leeming, David R Nisbet, Bryony A Nayagam

## Abstract

Cochlear implants partially restore sound sensation for individuals with severe-to-profound hearing loss. This can significantly improve the recipient’s quality of life, largely through improved communication with spoken language. However, surgical trauma from electrode insertion also instigates biological responses that can compromise device performance and longevity. To address these limitations, we designed a novel polymer-based coating that provides controlled release of selonsertib, an apoptosis signal–regulating kinase 1 (ASK1) inhibitor with well described anti-inflammatory, anti-fibrosis and cytoprotective properties. We describe the physical and molecular evaluation of this coating, including the application of next generation mass spectrometry–based proteomics in mammalian cochlear explants and their culture medium. These analyses provide new insights into the acute tissue response in the damaged cochlea and demonstrate that selonsertib eluted from the polymer coating has a significant pro-survival, anti-inflammatory, and anti-fibrosis effect. This study establishes proof-of-concept for selonsertib-eluting polymers, demonstrating both feasibility for targeted drug delivery and efficacy for mitigating acute cochlear damage responses.

## Introduction

The cochlear implant is a transformative medical device that can partially restore sound sensation for people with severe-to-profound hearing loss. Recipients report improved speech and communication abilities, with reduced depression, anxiety, and stress^1–4^. However, cochlear implant surgery can cause acute tissue damage within the cochlea, that initiates apoptosis and inflammation responses^5^. Subsequent immune cell infiltration drives the secretion of cytokines and reactive oxygen species that exacerbate cell death^5^. These immune infiltrates can migrate through cochlear lymph, damaging regions of the cochlea beyond the immediate site of electrode insertion^6^ and the vestibular organs^7^. Collectively, these innate responses can damage residual hearing and balance, reduce the population of neurons available for electrical stimulation, and prime the foreign body response^5^.

The foreign body response drives fibrotic tissue formation around 95-100% of cochlear implant electrodes^8,9^. This fibrotic encapsulation increases electrode impedance, necessitating stronger stimulation charges that degrade sound quality and impact device longevity. Commonly, moderate fibrosis causes *soft* cochlear implant failures, whilst severe fibrosis can contribute to migration or extrusion of the electrode^8–13^. Therefore, fibrosis represents a major limitation for cochlear implant function and protecting the electrode–tissue interface has emerged as an important research priority. From this research, slow-release electrode coatings have become established as an effective and clinically translatable method for drug delivery to the cochlea. To date, these coatings have primarily delivered neurotrophins or dexamethasone^5^. However, improved molecular profiling has expanded our understanding of the cochlear injury response, revealing new therapeutic targets that could concomitantly address the apoptosis, inflammation, and extracellular matrix remodelling that undermines cochlear implant electrode performance^5,14^. Among these, apoptosis signal-regulating kinase 1 (ASK1) is a compelling therapeutic target.

ASK1 is a highly conserved, ubiquitously expressed MAP3K that rapidly activates in response to cellular stress. As a central upstream regulator, ASK1 triggers diverse apoptosis and inflammation signalling cascades^15^. ASK1 activation occurs within minutes of ototoxic aminoglycosides being added to hair cells *in vitro*^14,16^ and ASK1 inhibition attenuates cell death in neomycin treated cochlear explants^16^. Additionally, ASK1 inhibition reduces apoptosis, inflammation, and fibrosis in a substantial number of surgical and disease models (reviewed in^15^). This success has led to the ASK1 inhibitor selonsertib progressing through phase I-III clinical trials, demonstrating human safety, tolerability, and efficacy^17–23^. Combined, these findings position selonsertib as an exciting candidate to mitigate the biological responses negatively impacting cochlear implant outcomes.

To evaluate selonsertib as an otoprotective agent, we developed a selonsertib-eluting coating for cochlear implant electrodes. Here, we describe the material characterisation and release kinetics of functional coatings that achieved sustained drug delivery. The biological activity of eluted selonsertib was subsequently confirmed using mass spectrometry-based proteomics, demonstrating a robust attenuation of damage responses in cochlear tissue. Collectively, these data establish a translationally actionable proof-of-concept for the selonsertib-eluting therapeutic strategy, whilst highlighting the analytical power of modern proteomics for pre-clinical discovery.

## Methods

### Polymer drug loading and coating

Polycaprolactone (PCL) and Polyethylene vinyl acetate (PEVA) were dissolved at 12% w/v in acetonitrile. For drug loaded coatings, a 40 μL solution containing selonsertib (100 μM) in DMSO was added to obtain a final concentration of 10% w/v polymer, and control coatings were prepared using 40 μL of DMSO in 12% w/v polymer to obtain a final concentration of 10% w/v for each polymer. Silicon filaments (0.4 mm diameter) mimicking electrodes and were coated with drug loaded or control PCL, or PEVA, via dip coating (1,3,5 or 10 layers each). Each coating condition was prepared in triplicate.

### Scanning electron microscopy

To assess coating morphology and thickness, polymer coated dummy electrodes were splutter coated using a Dynavac SC100 sputter coater at 25 mA for 40 seconds. Samples were imaged using a high-resolution Hitachi SU7000 scanning electron microscope (SEM) at 5 kV, with an 8 mm working distance.

### Selonsertib detection

Three-layer coatings were applied to dummy electrodes as described above. The coated dummy electrodes were then immersed in 1 mL of neurobasal cell culture media and kept at 37 °C under sterile conditions. Media was replaced hourly during the first eight hours, then daily for 28 days. High performance liquid chromatography (HPLC) was used to quantify selonsertib in collected medium, using gradient elution with a mobile phase (A) of 0.05% TFA in water and a mobile phase (B) of 100% ACN. The elution solvent increased from 5% to 95% over 8 minutes and was held at 95% B for 2 minutes (10 minutes in total). The column oven was held at 40 °C using a C18 column (5 μm) and the autosampler temperature was 20 °C. The sample injection volume was 5 μL and the mobile phase flow was 1.5 mL/minute. A calibration curve from 20 mM to 0.078 mM (10 calibration points) was prepared by dissolving selonsertib in DMSO and passing the sample through a 0.2 μm filter. Selonsertib was detected at a retention time (RT) of 5.7 minutes.

### Preparing Pre-conditioned Cell Culture Media

Three coatings of selonsertib loaded PCL or control PCL were applied to dummy electrodes as above. Coated dummy electrodes were then immersed in 1 mL of neurobasal cell culture medium and maintained under sterile conditions at 37 °C for 28 days. A media only control was also prepared, using submerged dummy electrodes with no coating applied. Conditioned media was stored at -20°C, before being defrosted on the day of cochlear explant collection. N2, L-glutamine, and glucose were added to the defrosted media immediately prior to being used for cell culture, per our published protocol^24^.

### Care and use of mice

In accordance with the 3Rs ethical framework, surplus pups from an unrelated study were used to minimise animal numbers. These pups were F1 BALB/c × C57BL/6 (CBB6 mice), bred using in-house colonies at The Florey Institute of Neuroscience and Mental Health (Animal Ethics Approval 19-100). Animal housing and research within this facility adheres to the National Health and Medical Research Council’s *Australian Code for the Care and Use of Animals for Scientific Purposes* (8th Edition 2013, 2021 update), the Victorian state Act for the Prevention of Cruelty to Animals 1986, and the Prevention of Cruelty to Animals Regulation 2008.

### Mouse Cochlear Explant Culture

P3 mouse pups were euthanised by cervical dislocation and immediately placed on wet ice. Cochlear explants were subsequently dissected and cultured as described in^24^, adapted slightly to facilitate proteomic analysis. Specifically, dissected explants were suspended in ice cold neurobasal media, until all dissections were complete. Explants were then moved to a low adhesion 96 well plate containing pre-conditioned media (150 μL). The free-floating explants were then incubated for 24 hours at 37 °C, 5% CO_2_.

After 24 hours culture, explants were aspirated into a wide bore pipette tip in 40 μL of media and gently ejected in 1 mL of room temperature PBS. This process was repeated two more times into fresh PBS, before the explant in 40 μL of PBS was moved into a 1.5 mL low bind Eppendorf tube containing 40 μL of 2x RIPA buffer (Millipore, 20-188) + protease inhibitor (Roche, cOmplete™ ULTRA EDTA free, 05892791001) and immediately frozen in liquid nitrogen and stored at -80°C. At the same time, 40 μL of culture media was collected from each well, and added directly into in a low bind Eppendorf tube containing 40 μL of 2x RIPA buffer + protease inhibitor and snap frozen for storage.

### Protein Extraction, Digestion, and Mass Spectrometry

Once all experimental replicates were performed, samples were simultaneously defrosted and moved to enclosed Covaris AFA strip tubes (TrendBio, 520292). Samples were then sonicated in the Covaris R230 ultra-sonicator, with the sonication program achieving an Average Incident Power of 112.5, using 60 iterations of 15 seconds on, 5 seconds rest at 6 °C (PIP 450, DF 25, CPB 1000, Dithering Enabled). In fresh low bind tubes, samples were centrifuged for 10 minutes at 1400 g to remove debris, and supernatant collected into new low bind tubes. After standard BCA quantification (Pierce BCA Protein Assay Kit, ThermoFisher, 23227), samples were normalised using ultrapure (Milli-Q) water and then processed using the standard S-trap micro protocol. In brief, 2x lysis buffer (10% SDS, 100 mM TEAB, Milli-Q water, pH 8.5) was added to each normalised sample at a 1:1 ratio and vortexed. 500 mM TCEP was added to a final concentration of 5 mM, incubated on a shaker (1000 rpm) at 37 °C for 45 minutes. 500 mM iodoacetamide was then added to a final concentration of 20 mM and incubated at room temp for 20 minutes in the dark. 28% phosphoric acid was then added, to achieve a minimum final concentration of 2.5%. Binding/wash buffer (100 mM TEAB in 90% methanol.) was added at 6X the total sample, and 170 μL of sample was then added to the s-trap column and centrifuged at 4000 g for 30 seconds. The remaining sample was added to the s-trap columns in batches of 170 μL and centrifuged until all sample had been processed. After additional wash steps, trypsin (dissolved in 50 mM TEAB) was added to the s-trap filter at a loading of 1 μg of trypsin per 10 μg protein, for overnight digestion at 37 °C in a humidified chamber. Protein was then eluted from the column filter using 3 elution buffers (50 mM TEAB in water, 0.2% formic acid in water, and 50% acetonitrile in water). Combined eluents were lyophilised and stored at -80°C. Resuspension in was performed in 0.2% formic acid in 2% acetonitrile just prior to mass spectrometry analysis.

### LC-MS/MS Analysis of Protein Digests

Peptide samples were analysed by LC-MS/MS using an Orbitrap Astral mass spectrometer coupled to a Vanquish Neo UHPLC (Thermo Fisher Scientific) using a trap-elute strategy. The trap and analytical columns were an Acclaim Pepmap nano-trap (Dionex—C18, 100 Å, 75 µm × 2 cm) and a 5.5 cm µPAC Neo respectively. The eluents were water with 0.1% *v/v* formic acid (solvent A) and 80% CH_3_CN with 0.1% *v/v* formic acid (solvent B). The flow rate was 750 nL/minute and the gradient was as follows [time (minute), %B]: [0, 3], [0.3, 6], [16, 23.5], [17.7, 40], [18.7, 50], [18.8, 99].

Column wash was then activated at 2 µL/min under combined flow and pressure control for 1.2 minutes followed by trap column wash (combined flow and pressure control with 2 zebra wash cycles at 80% CH_3_CN) and fast equilibration of the analytical column (combine flow and pressure control). Peptides were ionised via electrospray ionisation wherein the spray voltage was 1.9 kV and the ion transfer tube was held at 290 °C. The mass spectrometer was operated in the DIA mode. The MS1 scans were collected in the orbitrap analyser at resolution of 120,000 at *m/z* 200 over of range of 380-980 *m/z* in the profile mode. The MS1 AGC target was 500% with a maximum IT of 5 ms. MS2 DIA spectra were collected using the Astral mass analyser with isolation window of 2 *m/z*, normalized HCD energy of 27, normalised AGC target of 500% and maximum injection time of 3 ms. The cycle time was kept to 0.6 s.

### Proteomic Data Analysis

Raw mass spectrometry data were processed with Biognosys Spectronaut (v 20.1) using a directDIA strategy against reviewed Mouse protein sequences that were obtained from UniProt (August 2022). Post-translational acetylation of protein N-termini and carbamidomethylation of cysteine were set as fixed modifications and oxidation of methionine was configured as a variable modification. Peptides were quantified at the MS2 level. Precursor, peptide and protein false discovery rates were set at 0.1% and all other parameters were as default. The resultant protein abundance table was exported for subsequent analysis in R. Missing values were handled using singular value decomposition (SVD)-based imputation implemented in the pcaMethods package following log₂ transformation of intensities. Differential abundance analysis was performed using limma^25^. P-values were adjusted for multiple testing using the Benjamini–Hochberg false discovery rate (FDR) method. Proteins were considered differentially abundant at FDR < 0.05 and |log₂ fold-change| ≥ log₂(1.5). Gene set enrichment analysis (GSEA) was performed using clusterProfiler^26^ and ReactomePA^27^. Ranked gene lists (based on log₂ fold-change) were used for Gene Ontology (GO; Molecular Function, Cellular Component, Biological Process) and Reactome pathway enrichment analyses. Gene symbols were mapped to Entrez identifiers where required. Enrichment significance was assessed using FDR-adjusted P-values (Benjamini–Hochberg), with a cutoff of 0.05.

### Data Availability

Mass spectrometry proteomics data have been deposited to the ProteomeXchange Consortium (http://proteomecentral.proteomexchange.org) via The Proteomics Identifications Archive database (PRIDE)^28^ (https://www.ebi.ac.uk/pride/archive/), ID PXD074623 (Preprint note: data will be made open access once peer review is complete). Identified proteins and abundances are provided in text format in Supplementary Data 1.

## Results

### Polymer Encapsulation of Selonsertib

Polycaprolactone (PCL) and Polyethylenevinyl acetate (PEVA) were evaluated as selonsertib carriers. These polymers were selected based on their compatibility with selonsertib’s high hydrophobicity, as well as their flexibility, low cell attachment properties, and demonstrated biocompatibility in commercial biomedical applications^29^.

### Morphological Analysis of drug loaded coatings

To assess whether selonsertib-loaded PCL and PEVA could form structurally sound coatings, multiple layers were applied to dummy electrodes by dip coating. Up to 10 layers were successfully applied, with the first layers averaging ∼ 6-7 μm in thickness and subsequent layers increasing to ∼ 9-10 μm. Specific SEM measurements of total coating thickness for 1, 3, 5 and 10 layers were 6.5 μm ± 2 μm, 19.1 μm ± 1 μm, 45.5 μm ± 6 μm, and 92 μm ± 18 μm, respectively (n = 3 ± SD). Based on these dimensions, coatings of five layers or more were considered impractical for *in vivo* applications^30–32^. Thus, subsequent evaluations were performed using three layers of selonsertib-eluting polymer (Figure 1A).

**Figure 1.**
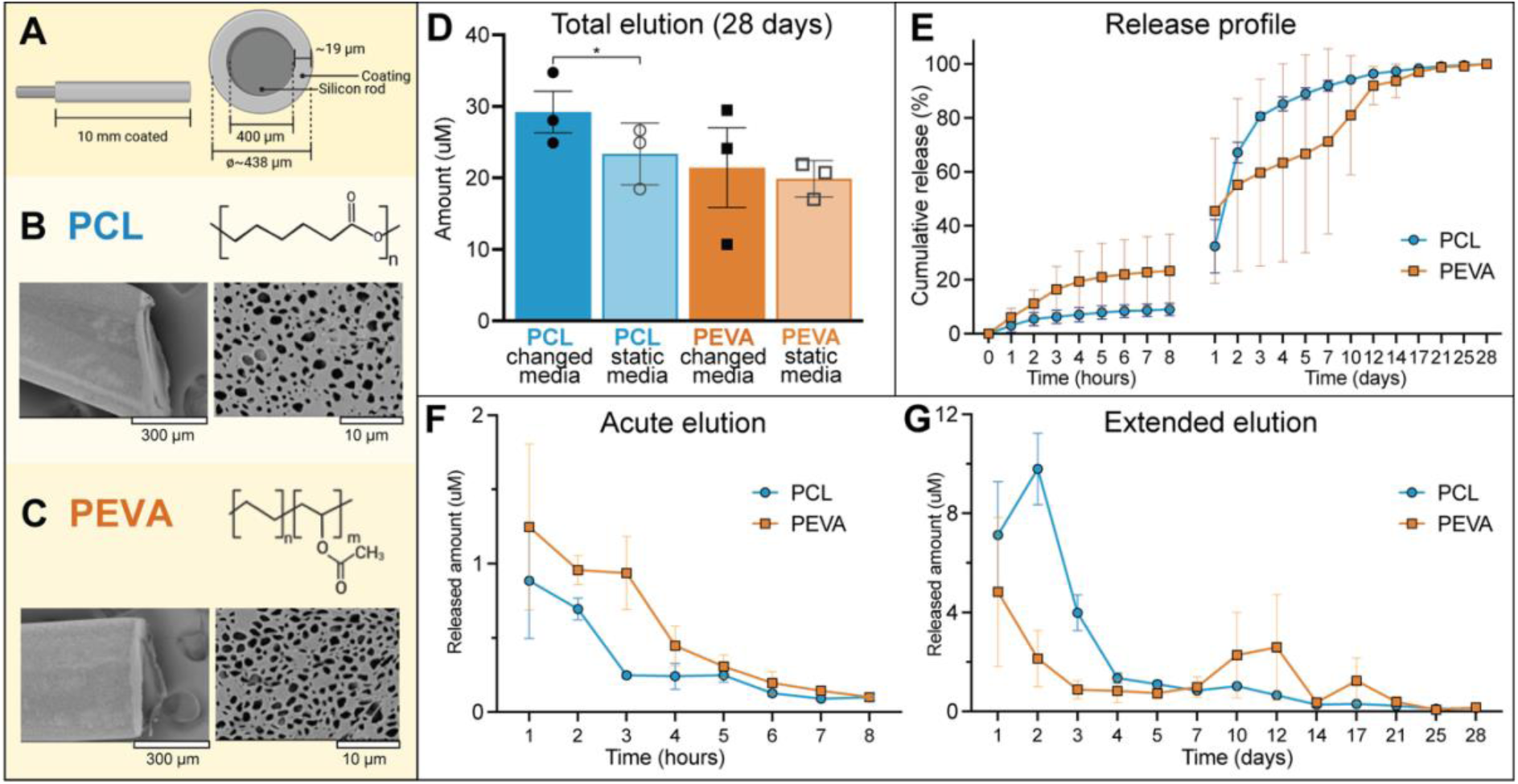
*Selonsertib* encapsulation and release. **A)** Three layers of PCL or PEVA containing 10% selonsertib were applied to dummy electrodes, producing a uniform coating with a porous surface. B) PCL chemical structure and representative SEM image of the selonsertib loaded PCL coating **C)** PEVA chemical structure and representative SEM image of the selonsertib loaded PEVA coating. **D)** Coated dummy electrodes were submerged in 1 mL neurobasal media for 28 days, which was analysed using HPLC to quantify selonsertib elution into the media. Two conditions were tested, a dynamic state, where media was changed daily (changed media) and a static state, where media was not changed for the 28-day experiment (static media). In both conditions, the PCL-based coating eluted significantly more selonsertib than the PEVA coating (*p < 0.05 based on one tailed t- test) **E)** Cumulative selonsertib release profiles for PCL and PEVA coatings, shown as the percentage of total selonsertib released over the 28-day experiment. Raw values for the changed media condition are given in **F) and G)** showing the quantity of selonsertib released into media over time. Media was replaced after each measurement during the first eight hours and then daily up to 28 days. All values calculated as the average of triplicates +/- SD.

SEM confirmed that the three layer selonsertib-eluting coating formed uniform, porous surfaces on dummy electrodes (Figure 1B,1C). The average pore diameter of PCL (2126.5 ± 1162 μm) was slightly larger than PEVA (1809.5 ± 1607 μm). However, the difference in pore size was not statistically significant (p > 0.79, two tailed t-test).

### Selonsertib release from PCL and PEVA

Coated dummy electrodes were submerged in 1 mL of neurobasal media at 37 °C. High-performance liquid chromatography (HPLC) was then used to detect selonsertib that was released into the media (Figure 1D-G). Two conditions were tested, a static state with no media changes, and a dynamic state, where complete media changes were performed daily to mimic cochlear fluid turnover. Under both conditions, PCL released more selonsertib than PEVA (Figure 1D), averaging ∼1 μM per day over the 28-day sampling period, compared to 0.78 μM per day for PEVA. However, daily measurements revealed more variability (Figure 1E–G), with PCL releasing approximately 5% of the total eluted drug within the first 24 h (∼7 μM), followed by 10 μM at 48 h, 4 μM at 72 h, and stabilising at ∼1 μM for the remainder of the first week. In contrast, selonsertib-eluting PEVA had a drug burst after one week, with notable variability observed between replicates. For both polymers, less selonsertib was released in the static condition when compared to the daily media change. This trend was statistically significant for PCL, but not for PEVA (p > 0.4, one tailed t-test) likely due to the higher variability observed in PEVA-selonsertib release.

As selonsertib’s EC_50_ in humans is reported at 56 ng/mL (∼0.1 μM in whole blood)^18^, both PCL and PEVA achieved therapeutically relevant selonsertib release profiles. However, in the context of cochlear implantation, where acute apoptosis and inflammation is induced by electrode insertion, we considered PCL’s initial drug release to be beneficial. Thus, for subsequent *in vitro* proteomics, selonsertib-eluting PCL was used.

### The molecular effect of PCL and selonsertib

Cochlear explants were used to model the tissue damage response initiated during cochlear implant electrode insertion. Cochlear explants provide a robust *ex vivo* model, with excellent cell survival during short-term culture^16,24^. However, dissection disrupts the cochlear microenvironment and shears peripheral margins of the neurosensory epithelium^24^. We hypothesised that this disruption would trigger acute apoptosis and inflammatory signalling, and that selonsertib could attenuate these responses.

To test these hypotheses, we pre-conditioned basal media for 28 days at 37 °C, containing either a submerged selonsertib-eluting PCL coated electrode, a PCL coated electrode (vehicle control), or an uncoated electrode (no coating-media control). This pre-conditioned media was then used to culture cochlear explants for 24 hours. Subsequent proteomic analysis detected ∼7,700 protein groups in each explant preparation. Culture media was also collected from each explant to evaluate secreted chemotactic signals, with ∼1,700 proteins detected. Summary data are presented below (Figures 2-4) and complete protein results are provided in Supplementary Data 1.

**Figure 2.**
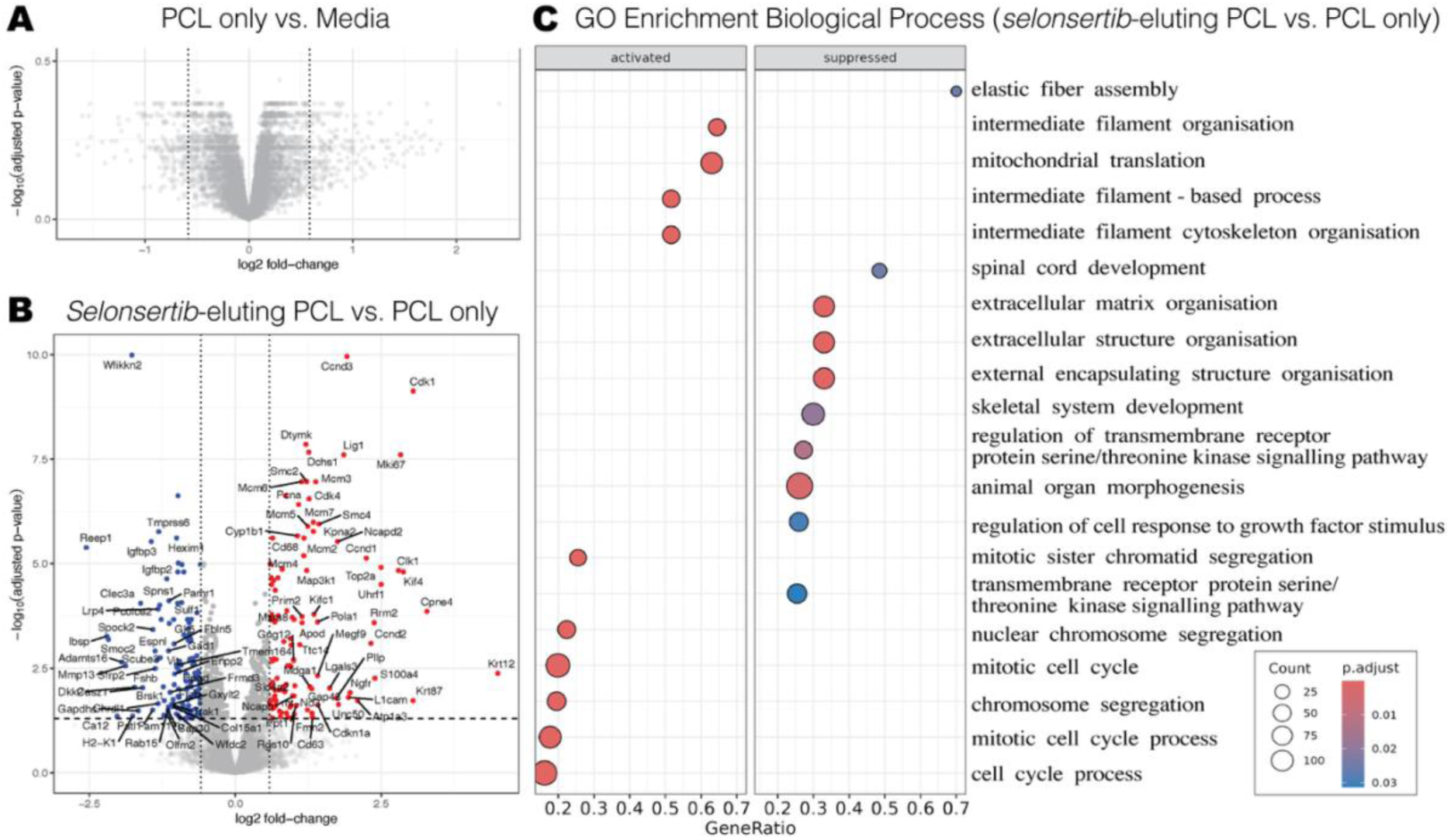
Selonsertib suppresses the acute damage response in cochlear explants. **A)** Volcano plot showing no significantly different protein expression between cochlear explants cultured in PCL pre-conditioned media (PCL only) and those cultured in the uncoated electrode pre-conditioned media (Media) (n = 7 explants per group, FDR < 0.05). **B)** Volcano plot showing different protein expression between the cochlear explants cultured in selonsertib-eluting PCL pre-conditioned media and PCL only pre-conditioned media (n = 7 explants per group). Significantly upregulated proteins are represented in red on the right, and significantly downregulated proteins are blue on the left. Protein groups with no significant changes are shown in grey. Significance was defined as a ≥ 1.5-fold change in protein abundance and an FDR > 0.05 (significance thresholds are indicated by dashed lines on each plot). A complete protein list is provided in Supplementary Data 1. **C)** Gene set enrichment analysis of protein abundances ranked by fold-change (selonsertib-eluting PCL compared to PCL only control) against GO biological processes. Dot size reflects the number of quantified proteins associated with each GO term and dot colour indicates the p-value adjusted for multiple comparisons. The proteins reduced in the selonsertib-eluting PCL group are represented as suppressed, whilst increased protein number is represented as activated.

**Figure 3.**
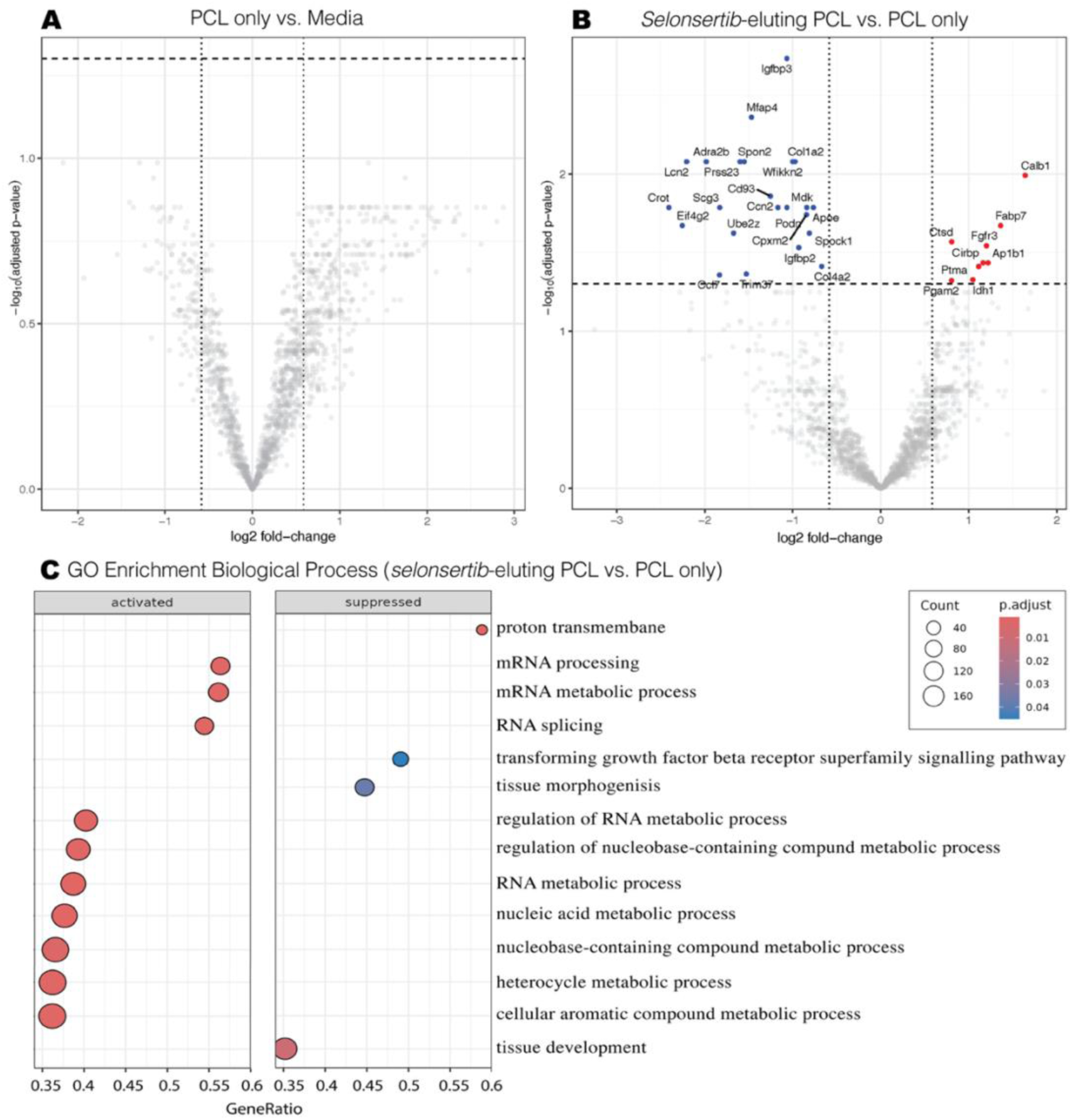
Key secreted inflammation signals are reduced in the culture medium of selonsertib treated cochlear explants. **A)** Volcano plot demonstrating no differentially expressed proteins (DEPs) when comparing culture media collected from cochlear explants, after 24 hours culture in either PCL pre-conditioned media or an uncoated-media control (n = 7 explants per group, FDR < 0.05). **B)** Volcano plot showing DEPs detected when comparing the media collected from cochlear explants cultured in selonsertib-eluting PCL pre-conditioned media and those cultured in PCL only pre-conditioned media (n = 7 per group). Significantly upregulated proteins are represented in red on the right, and significantly downregulated proteins are blue on the left. Protein groups with no significant changes are shown in grey. Significance was defined as a ≥ 1.5-fold change in protein abundance and an FDR > 0.05 (these thresholds are indicated by dashed lines on each plot). **C)** Gene set enrichment analysis of protein abundances ranked by fold-change (selonsertib-eluting PCL compared to PCL only control) against GO biological processes. Dot size reflects the number of proteins associated with each GO term and dot colour indicates the p-value adjusted for multiple comparisons. Proteins reduced in the selonsertib-eluting PCL group are represented as suppressed, whilst increased protein number is represented as activated.

**Figure 4.**
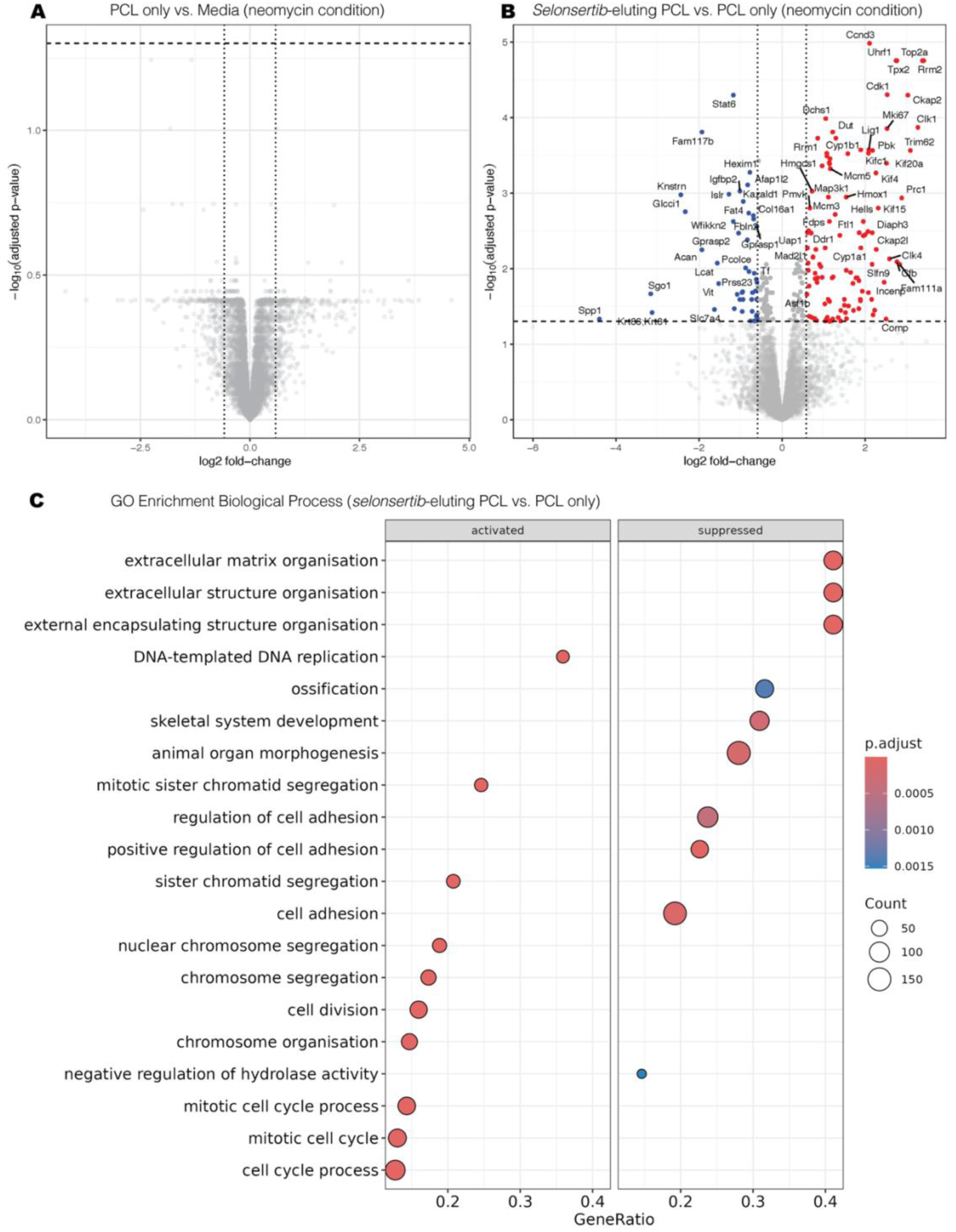
Selonsertib reduces inflammation, apoptosis and fibrosis associated proteins in neomycin treated cochlear explants. **A)** Volcano plot showing no DEPs when comparing cochlear explants after 24 hours of culture in PCL pre-conditioned media (PCL only) or uncoated electrode pre-conditioned media (Media) (both supplemented with 1 mM neomycin). Media (control) n = 4, PCL only (vehicle control) n = 3, selonsertib-eluting PCL n = 4. **B)** Volcano plot showing DEPs detected when comparing cochlear explants cultured in selonsertib-eluting PCL pre-conditioned media and those cultured in PCL only pre-conditioned media. Significantly upregulated proteins are represented in red on the right, and significantly downregulated proteins are blue on the left. Protein groups with no significant changes are shown in grey. Significance was defined as a ≥ 1.5-fold change in protein abundance and an FDR > 0.05 (these thresholds are indicated by dashed lines on each plot). C) Gene set enrichment analysis of protein abundances ranked by fold-change (selonsertib-eluting PCL compared to PCL only control) against GO biological processes. Dot plot representing proteins significantly altered by selonsertib-eluting PCL when compared to PCL only control, using GO enrichment terms categorised by Biological Processes. Dot size reflects the number of proteins associated with each GO term and dot colour indicates the p-value (adjusted for multiple comparisons). Proteins reduced in the selonsertib-eluting PCL group are represented as suppressed, whilst increased protein number is represented as activated.

### PCL-conditioned media has no acute molecular effect on cochlear explants

Because polymers can leach byproducts that induce immune reactions, we compared cochlear explants cultured in PCL-pre-conditioned media with those cultured in no-coating control media. No significant difference in protein expression was observed between the groups (Figure 2A, FDR < 0.05) Consistent with these findings, analysis of secreted proteins in post-culture media showed no difference between the PCL-pre-conditioned and control media cultured explants (Figure 3A, FDR < 0.05).This finding was replicated in independent experiments, internal replicates (Supplementary Data 2) and when an additional stressor (neomycin) was added (Figure 4A). These data indicate that PCL did not release byproducts capable of eliciting an acute molecular response.

### Selonsertib reduces inflammation and fibrosis-associated protein expression in cochlear explants

To evaluate the biological activity of selonsertib, we compared the proteomic profile of cochlear explants that had been cultured in selonsertib-eluting PCL pre-conditioned media with those exposed to PCL-only pre-conditioned media. This analysis identified 115 significantly upregulated proteins and 129 downregulated, confirming that selonsertib retained chemical activity following polymer encapsulation and prolonged aqueous liquid submersion (Figure 2B, Supplementary Data 1).

Upregulated proteins were predominantly associated with metabolism, proliferation, debris clearance, and repair (Figure 2B, C). A notable number of anti-inflammatory proteins were also upregulated, including RGS10^33^, ABCA1^34^, TMEM176B^35^, CBR2^36^, GNG12^37^ and GRNC^38^. Furthermore, the anti-oxidant DHCR24^39^ was upregulated, alongside pro-survival proteins MAP3K1^40,41^, NDUFA4^42^, MCL1^43^, MT-ND3^44^, TIMM8B^45^, LITAF^46^, and LAMP2^47^. In contrast, a single apoptosis associated protein (NRADD^48^) was upregulated. Collectively, these proteomic signatures are consistent with reduced inflammation, enhanced mitochondrial function, and DNA damage repair characteristic of an active pro-survival response.

A complementary pro-survival signal was also evident among downregulated proteins, including reduced expression of pro-apoptosis DKK2^49^, DKK3^50^, and HEXIM1^51^, as well as suppression of the inflammation associated ENPP2^52^, STAT6^53^, NFAT^54^, and ARG2^55^. Most striking, was the reduction of ECM/fibrosis associated proteins, including IGFBP2^56,57^, CCN2^5^, LOXL2^58^, SFRP1, SFRP2^5,59^, IBSP^60^, SMOC2^61^ and many more (Supplementary Data 1). Several of these proteins are part of the WNT signalling pathway, indicating that selonsertib interrupts the self-sustaining, fibrosis enhancing WNT-TGF-β feedback loop. This observation is further supported by the reduced expression of TGF-β companion, chaperone, and activator proteins (ADAMTS16^62,63^, WFIKKN2^64^, THBS1^65^, and LTBPS^66^).

### Selonsertib suppresses secretion of inflammation recruitment and ECM deposition signals

To evaluate the secretome of cochlear explants in the experiments above, we also collected culture media for proteomic analysis. In total, 27 differentially expressed proteins (DEPs) were detected, with the majority downregulated. Only 9 DEPs were upregulated by selonsertib, which were preponderantly intracellular metabolism-associated proteins (Figure 3C). These data may reflect stress induced cell leakage from a greater number of surviving cells. However, non-adherent cells in the media may also contribute to this signal. In contrast, 18 (DEPs) were downregulated, with well-defined immune recruitment and pro-ECM properties. These DEPs include PRSS23^67^, SPON2^68^, MFAP4^69^, MDK^70^, SPOCK1^71^, COL1A2, COL4A2^72,73^, and IGFBP 3 (Figure 3). IGFBP3 is a well characterised activator of ECM deposition, apoptosis^74^, and inflammation^75^, whilst IGFBP2 has been primarily documented as a proliferation signal for malignant cancers. However, there is a growing body of evidence that IGFBP2 is also an activator of ECM and apoptosis responses, which is supported by these data in a cell stress model^56,76^.

GO enrichment analysis highlighted a coordinated reduction of TGF-β superfamily signalling (Figure 3C) with most of the selonsertib-downregulated proteins being downstream responders of TGF-β. This corresponds with the likely interruption of the WNT-TGF-β pathway identified within the cochlear explant cells above. However, our analysis also detected significantly reduced WFIKKN2 in both cochlear explants and their secretome. As WFIKKN2 is a highly potent antagonist of GDF8/11 and related TGF-β-family ligands, its reduced secretion is likely to further impact the TGF-β fibrotic response^64,77^. Given that TGF-β is a master regulator of fibrosis throughout the body, the suppression of TGF-β signalling observed in the selonsertib-treated explants and their secretome is a significant molecular outcome^5^.

### Selonsertib treatment remains protective during intensified cochlear injury

In the dissection-only conditions above, selonsertib significantly suppressed key inflammation, apoptosis, and extracellular matrix responses in cochlear explants. These data align with observations from other surgical and disease models^15^. However, ASK1 inhibition with a selonsertib analogue has also been shown to reduce hair-cell loss in aminoglycoside-treated cochlear explants^16^. Therefore, we used aminoglycoside exposure as a complementary, severe stress model, to further interrogate the anti-apoptotic effect ASK1 inhibition. Explants were dissected and cultured as above, with 1 mM of ototoxic neomycin included in the culture media.

Selonsertib significantly downregulated 46 proteins in the neomycin condition, which is fewer than in the dissection only selonsertib group. This difference could indicate a stronger damage response induced by neomycin but may also be caused by the smaller sample size of the neomycin cohort. Notably, 16 proteins were consistently downregulated by selonsertib when comparing the dissection-only and neomycin-treated explants, including pro-apoptosis HEXIM1^51^, inflammation associated STAT6^53^, COL16A1^78^, LAMB3^79^ and ZFP36L1^80^, ECM/fibrosis associated IGFBP2^56,57^, WFIKKN2^64^, LRP4^81^, PCOLCE^82^ SLIT2^83^, SPOCK2^84^, Epithelial–Mesenchymal Transition protein ARHGAP18^85^ and the less well-defined FAM117B, ISLR, FAT4^86^ PCOLCE^82^, and VITRIN^87^. This shared protein signature highlights that cochlear explant dissection itself elicits a core inflammatory and fibrotic response that is mitigated by selonsertib treatment.

Interestingly, the downregulated proteins that were unique to the neomycin + selonsertib group continued a pattern of reduced ECM signalling, including SPP1^88^, ACAN^81^, ITGBL1^89^, FBLN2^90^ and CNTNAP4^91^; with immune recruitment or activation also attenuated via downregulated QPCT^92^, VCAM1^93^, STAT5A^94^, MPEG1^95^ and PLA2G7^96^. As predicted, additional apoptosis associated proteins were supressed by selonsertib in the neomycin treated explants, including TF, a marker of Ferroptosis^97^ and ENDOG, a well-defined mitochondrial nuclease involved in apoptosis/autophagy^98^. Together, these data demonstrate that ASK1 inhibition not only suppresses inflammatory and ECM-associated responses but also attenuates diverse apoptotic pathways, providing multifaceted cell protection.

Among the 108 proteins that were upregulated by selonsertib, a substantial overlap (∼40% of DEPs) was again observed between the dissection-only and neomycin-treated explants. These included hair cell survival associated MAP3K1 and CDK1^40,41,99,100^ and several proteins associated with cell cycle processes, such as metabolism and proliferation (Figure 4C). Among the upregulated proteins that were unique to the selonsertib + neomycin group, were the cytoprotective HMOX1^101^ and FTL1^102^, which are well established antioxidants in the oxidative injury response. The upregulation of these stress-adapting proteins in the ototoxicity model reiterates that selonsertib can broadly attenuate cell-death pathways, whether triggered by well-defined redox signalling arising from aminoglycoside-induced mitochondrial damage, or by the more heterogeneous signals associated with physical trauma and inflammation.

Interestingly, despite the broad reduction of ECM–associated proteins following selonsertib treatment, DDR1 expression was also increased. DDR1^103^ is a structural collagen that contributes to anchoring of the basilar membrane and lateral wall, maintains tension and elasticity of the organ of Corti, and maintains the motile cytoskeleton of outer hair cells. The observed increase in DDR1 may reflect a compensatory response preserving mechanical integrity under stress. However, in the context of ototoxicity, enhanced outer hair cell survival due to ASK1 inhibition^16^ provides another possible explanation for elevated DDR1 levels.

## Discussion

ASK1 inhibition has emerged as a promising therapeutic strategy for preventing cell death, inflammation, and fibrosis, which are common biological responses known to reduce cochlear implant efficacy. Therefore, we hypothesised that the ATP-competitive ASK1 inhibitor selonsertib (IC50 8.1 μM) could attenuate biological responses in damaged cochlear tissue. Notably, selonsertib is well tolerated in humans and has an excellent oral bioavailability^15^. However, *first-pass effects* would likely reduce systemic availability and the blood-labyrinth barrier could further impact drug delivery to the ear^104^. In contrast, a drug eluting implant coating would directly release selonsertib into the cochlea. Therefore, we evaluated candidate drug carriers to adhere to implant electrodes and facilitate prolonged selonsertib elution. We performed material and molecular analyses, demonstrating the feasibility and efficacy of this novel therapeutic strategy for attenuating innate cochlear damage responses.

### Polymer Encapsulation Enables Sustained Selonsertib Release

Our material characterisation of selonsertib eluting PCL and PEVA confirmed that both polymers formed uniform, porous layers on dummy electrodes. This is consistent with previous reports that single dip PCL coatings do not impact electrode smoothness or flexibility^105^. However, our data builds on this, demonstrating that both PCL and PEVA can be layered on electrode surfaces, raising the potential for prolonged, timed, or combination drug delivery. Our data also demonstrated that both polymers achieved prolonged selonsertib release over a 28-day period, at concentrations above the reported human EC50 of ∼ 0.1 µM^18^.

Interestingly, less selonsertib was released in the static media condition, suggesting that drug accumulation may reduce diffusion. This effect could prolong selonsertib release *in vivo*, although it seems unlikely given the fluid turnover that occurs within the cochlea. We were unable to test this *in vitro*, as modelling cochlear fluid dynamics remains a key challenge for preclinical testing of inner ear therapeutics. Microfluidic platforms combined with 3D printing may eventually enable directional flow models *in vitro*. However, the complexity of outer hair cell stirring^106^, blood labyrinth barrier leakage^107^, CSF perilymph interchange^108^, and other unknown cellular interactions cannot be adequately replicated *ex vivo*. Therefore, animal models remain essential for subsequent efficacy testing, despite anatomic differences between species^30^. In our *in vitro* model, we accounted for fluid turnover by using a daily media change. This condition included a volume 20 times larger than that of the human cochlea^30^ which may have resulted in a conservative estimate of elution efficacy. Nevertheless, it provides greater confidence that the observed drug elution and associated molecular effects will not be diluted when tested *in vivo*.

### Proteomics Reveals Acute Biological Signals

To evaluate the molecular effect of the novel selonsertib-eluting coating, we defined the proteomic landscape of murine cochlear explants and their secretome using quantitative mass-spectrometry. This proteomic analysis revealed that a robust damage response occurs within 24 hours of dissection, likely due to the disruption of the cochlear microenvironment and tissue shearing during collection. However, when explants were cultured in selonsertib-eluting PCL pre-conditioned media, these responses were profoundly attenuated. These data confirm that selonsertib remained chemically active, even after the encapsulation process and subsequent elution at 37 °C. In contrast, no changes were detected in response to PCL-only-conditioned media, consistent with *in vivo* studies that infer longer term PCL biocompatibility^105^.

The selonsertib-suppressed proteins in cochlear explants aligned unambiguously with reduced apoptosis and inflammation signalling, and the suppression of extracellular matrix formation. Several DEPs indicated that the WNT-TGF-β-SMAD fibrotic feedback loop^5^ had been interrupted, which was supported by a complementary suppression of TGF-β-associated proteins in the explant secretome. Notably, explants were cultured in a volume of media larger than that of the human cochlea^30^. Therefore, it is possible that the detected molecular response is somewhat underestimated. However, the detection of secretome changes in a four-fold dilute volume, highlights both the magnitude of the secreted signal and the analytical sensitivity of high-resolution mass spectrometry.

In parallel to the suppression of apoptosis, inflammation, and fibrosis signals resulting from selonsertib treatment, several pro-survival and repair-associated proteins were upregulated. A similar pro-survival signal was also observed when cellular damage was intensified using neomycin. However, in the ototoxic condition, additional antioxidant and stress-adaptive proteins were also upregulated in the selonsertib treatment group. These data show that selonsertib induces a molecular shift away from acute damage-responses to controlled recovery, even when activated by different cell stressors. This remarkable impact was further demonstrated by the concomitant reduction of proteins that have been independently implicated in fibrotic pathology and evaluated as a stand-alone therapeutic targets (DHCR24^39^, MFAP4^69^, SFRPs^59,109–111^, SMOC2^61^, THBS1^112^, ADAMTS1^62,63^, ENPP2^52^, and IGFBP2^56^). Selonsertib’s simultaneous attenuation of these targets reinforces the notable advantage of inhibiting an upstream regulator of highly conserved signalling cascades^15^.

Another notable signal to arise in the selonsertib treated explants, was the upregulation of cell cycle proteins. Initially, we hypothesised that this was due to selonsertib preserving cellular homeostasis by dampening stress-activated apoptosis and inflammatory pathways^16^. Indeed, early cell-cycle arrest is a hallmark of stress-induced apoptosis, and our selonsertib dataset showed suppression of apoptosis associated proteins. However, a recent molecular docking study identified that selonsertib also binds to CDK6 (IC50 9.8 μM)^113^. Thus, the widely reported reduction of apoptosis, inflammation, and fibrosis associated with selonsertib, could be due to a combined modulation of stress-activated and cell-cycle–associated pathways. For example, ASK1 and CDK6^100^ inhibition have been used independently to attenuate inflammation responses in the 5xfad mouse model of Alzheimer’s disease^15,114^. Unfortunately, this may also mean that the upregulation of cell cycle proteins in our data set was not a direct response to reduced apoptosis, but rather compensatory mechanisms activating in response to selonsertib inhibition of CDK6. Whilst the extent to which ASK1 and CDK6 inhibition contributes to selonsertib’s ability to reduce apoptosis, inflammation, and fibrosis across disease models warrants further investigation, the beneficial outcomes of selonsertib treatment are robust and replicated^15^. Furthermore, this unexpected dual target creates a new therapeutic opportunity, as CDK6 inhibitors are also effective against hormone receptor–positive breast cancer. In this respect, selonsertib could be used to reduce tumour growth whilst protecting against ototoxic outcomes in combined oncology treatments^115^. Certainly, several proteins associated with tumour metastasis were downregulated by selonsertib in our proteomics data, and we have previously described the otoprotective effect of ASK1 inhibition for aminoglycoside ototoxicity^16^. Together, these findings highlight an intriguing possibility for selonsertib repurposing in oncology.

Whilst the proteomic analyses in this study focused on the therapeutic effect of selonsertib, it also generated large datasets useful for other aspects of inner ear research. For example, KRT12 is reported to be specifically expressed in corneal epithelial cells of the eye. However, our analysis identified KRT12 to be one of the most differentially expressed proteins in the selonsertib vs vehicle comparisons. This provides protein-level validation of previous transcriptomic analyses that identified *Krt12* in mouse hair cells^116,117^, the zebrafish neuromast^118,119^ and as a dynamically changing signal during otic development in the frog^120^. Combined, these data suggest an underappreciated role for KRT12 in the structural maintenance of sensory epithelia. This example represents only one of the 7000+ proteins that were detected in each cochlear explant, which is a proteomic depth two to seven-fold higher than previous inner ear focused proteomics studies^121–125^.

This improvement is largely attributable to the Orbitrap Astral mass spectrometer, that quantifies up to five times more peptides than previous instrumentation^126^. However, bulk sonication in a temperature-controlled, non-contact format also significantly improved sample recovery and reproducibility. These advances underscore the growing utility of next generation proteomics for inner ear discovery.

### Selonsertib for Inner Ear Protection and Beyond

In summary, this study provides proof-of-concept for selonsertib-eluting polymers, demonstrating feasibility for drug delivery, and efficacy for mitigating acute cochlear damage responses. We initially predicted that selonsertib would predominantly affect cellular apoptosis and inflammation signals, due to the short experimental timeframe in which *ex vivo* cochlear explants can be maintained. However, proteomic analysis revealed that in addition to attenuated apoptosis and inflammation signals, a pronounced suppression of the extracellular matrix response also occurred. This broad suppression of stress-induced biological responses positions selonsertib as a strong therapeutic candidate for preserving cochlear function and for mitigating the fibrotic encapsulation of implanted devices.

For cochlear implant electrodes, moderate fibrosis diminishes sound quality, whilst severe fibrosis can lead to complete implant failure^8–13,127–130^. In these cases, where revision surgery is required, fibrosis levels are a primary predictor of revision success^12^. Therefore, even a modest reduction of fibrosis could have substantial clinical impact, improving audiological outcomes for cochlear implant recipients and extending the lifespan of their device. Selonsertib’s concomitant mitigation of apoptosis and inflammation following electrode insertion would also likely improve audiologic outcomes and has the potential to enhance recipient recovery. Indeed, further preclinical studies assessing long-term functional and electrophysiological outcomes using selonsertib eluting coatings are now warranted. However, given the established biocompatibility of PCL in the ear^105^ and the availability of human clinical safety data for selonsertib^15^, this strategy is positioned as an imminently translatable approach for improving implanted device outcomes. Furthermore, as the selonsertib-eluting coating could be applied to other implantable biomedical devices, there is strong potential for therapeutic expansion to neural electrodes, cardiac devices, ophthalmic implants, and other devices impacted by fibrotic encapsulation.

## Acknowledgements

We gratefully acknowledge the technical expertise and support provided by the staff at The Florey Bioservices Facility and the Melbourne Mass Spectrometry and Proteomics Facility (Bio21 Molecular Science and Biotechnology Institute). We also thank Maria Tanzer and Kael Schoffer (WEHI) for generously sharing equipment, and Marina Skrypnik for enabling this connection. This research was funded by The Passe and Williams Memorial Foundation Junior Fellowship (JMO) and Senior Fellowship (BAN), the MDHS Innovation Seed Grant, Early Career Researcher Grant, Graeme Clarke Seed fund, and Dept of Audiology and Speech Pathology, University of Melbourne (JMO). L.M.C.A was supported by an NHMRC Investigator Grant (GNT2041943) and DRN was supported by an NHMRC Ideas Grant (GNT1144996), and ARC Future Fellowship and Discovery Grant (DP220102549).

## Declaration of generative AI and AI-assisted technologies in the manuscript preparation process

ChatGPT (OpenAI, 2025–2026) was used for minor language editing during early manuscript preparation. After using this tool, the authors reviewed and edited the content extensively, and take full responsibility for the content of the published article.

