## Supplementary Data 2 for "Selonsertib-Eluting Electrode Coating Attenuates Cochlear Injury Pathways"

To evaluate the effect of selonsertib, we compared the selonsertib-eluting PCL and PCL only (vehicle) conditions. This comparison yielded results extremely similar to the selonsertib-eluting PCL vs. uncoated control (media) comparison, with ~90% of differentially expressed proteins (DEPs) overlapping. This overlap was 100% when considering the top 10 DEPs, reinforcing that PCL has no measurable effect on basal explant media composition. These data provide further evidence that PCL exposure does not have an acute molecular effect in cochlear explants, whilst serving as an internal replicate, increasing our sample size and demonstrating the reproducibility of our protein detection pipeline.

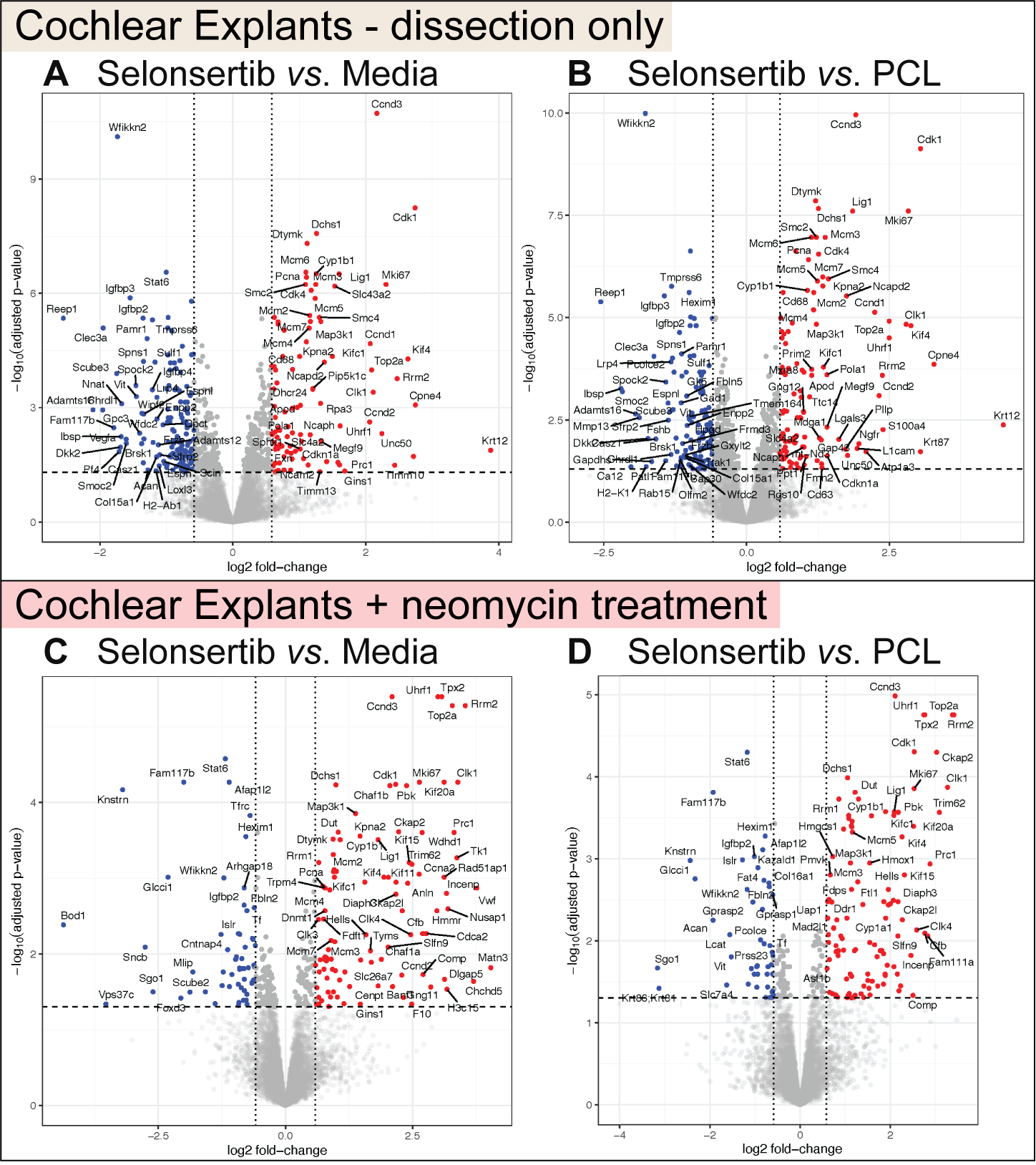

**Supplementary Figure 2.** Volcano plots showing up and downregulated proteins, when comparing the explants cultured in selonsertib-eluting-PCL preconditioned media, versus PCL only or the uncoated media control. **A and B** show data from explants cultured in preconditioned media only, whilst **C and D** are from a separate experiment in which preconditioned media was supplemented with neomycin to induce further damage. Panel B is a reproduction of Figure 2B in the main text, and D is a reproduction of 4B.
